# Repurposing Remdesivir for COVID-19: Computational Drug Design Targeting SARS-CoV-2 RNA Polymerase and Main Protease using Molecular Dynamics Approach

**DOI:** 10.1101/2023.06.15.545129

**Authors:** Mita Shikder, Kazi Ahsan Ahmed, Abu Tayab Moin, Rajesh B. Patil, Tasnin Al Hasib, Mohammad Imran Hossan, Deera Mahasin, Mohammad Najmul Sakib, Iqrar Ahmed, Harun Patel, Afrin Sultana Chowdhury

**Affiliations:** Department of Biochemistry and Molecular Biology, Bangladesh Agricultural University, Mymensingh, Bangladesh; Department of Genetic Engineering and Biotechnology, Faculty of Biological Sciences, University of Chittagong, Chattogram, Bangladesh; Sinhgad Technical Education Society’s, Sinhgad College of Pharmacy, Department of Pharmaceutical Chemistry, Off Sinhgad Road, Vadgaon (Bk), Pune 411041, Maharashtra, India; Department of Biochemistry and Molecular Biology, Bangabandhu Sheikh Mujibur Rahman Science and Technology University, Gopalganj, Bangladesh; Department of Biotechnology and Genetic Engineering, Noakhali Science and Technology University, Noakhali, Bangladesh; Biotechnology Program, Department of Mathematics and Natural Sciences, School of Data and Sciences, BRAC University, Dhaka, Bangladesh; Division of Computer-Aided Drug Design, Department of Pharmaceutical Chemistry, R. C. Patel Institute of Pharmaceutical Education and Research, Maharashtra, India

**Keywords:** SARS-CoV-2, RNA-dependent RNA polymerase (RdRp), Main protease (MPro), Remdesivir, Modified derivatives, Molecular docking, Molecular dynamics simulation.

## Abstract

The coronavirus disease of 2019 (COVID-19) is a highly contagious respiratory illness that has become a global health crisis with new variants, an unprecedented number of infections, and deaths and demands urgent manufacturing of potent therapeutics. Despite the success of vaccination campaigns around the globe, there is no particular therapeutics approved to date for efficiently treating infected individuals. Repositioning or repurposing previously effective antivirals against RNA viruses to treat COVID-19 patients is a feasible option. Remdesivir is a broad-spectrum antiviral drug that the Food and Drug Administration (FDA) licenses for treating COVID-19 patients who are critically ill patients. Remdesivir’s low efficacy, which has been shown in some clinical trials, possible adverse effects, and dose-related toxicities are issues with its use in clinical use. Our study aimed to design potent derivatives of remdesivir through the functional group modification of the parent drug targeting RNA-dependent RNA polymerase (RdRp) and main protease (MPro) of SARS-CoV-2. The efficacy and stability of the proposed derivatives were assessed by molecular docking and extended molecular dynamics simulation analyses. Furthermore, the pharmacokinetic activity was measured to ensure the safety and drug potential of the designed derivatives. The derivatives were non-carcinogenic, chemically reactive, highly interactive, and stable with the target proteins. D-CF3 is one of the designed derivatives that finally showed stronger interaction than the parent drug, according to the docking and dynamics simulation analyses, with both target proteins. However, *in vitro* and *in vivo* investigations are guaranteed to validate the findings in the future.

## 1. Introduction

Our world has witnessed a varied spread of previously unknown coronaviruses during this century, facilitated by rapid urbanization and ecological alteration of vulnerable public health structures [1]. In late December 2019, the coronavirus disease 2019 (COVID-19) pandemic began in Wuhan. From there, it has spread rapidly to more than 230 countries [2] at a rate beyond imagination, rampaging the world and becoming a global public health crisis. As of writing, there are 263,563,622 confirmed cases of COVID-19 resulting in 5232562 deaths worldwide [3]. Severe Acute Respiratory Syndrome Coronavirus-2 (SARS CoV-2), the causative agent of COVID-19, can infect both animals and humans with mild to severe respiratory, hepatic, and gastrointestinal complications [4]. Clinical data show that COVID-19 patients experienced various lethal consequences, including severe respiratory sickness, multi-system organ failure, and death. Additionally, it is clear from reports that older patients and those who already have respiratory or cardiovascular conditions are most at risk for infection [5]. SARS-CoV-2 could transmit through saliva, droplets, or secretions from an infected person’s nose after coughing, sneezing, and yawning, even while speaking, according to transmission pattern analysis [6].

The nucleocapsid core of the SARS CoV-2 contains a spike (S) protein, a membrane (M) protein, and an envelope (E) protein. The nucleocapsid core also contains a positive-sense single-stranded RNA genome (30 kilobases, kb) [7]. The nucleocapsid (N) protein, which encodes 4 structural proteins and 16 non-structural proteins (NSP), packages the virus’s RNA into a helical nucleocapsid [8–9]. Because it is an RNA virus, SARS-CoV-2 may create the versatile enzyme RNA-dependent RNA polymerase (RdRp), which is necessary for genome replication and transcription [10]. On the other hand, the main protease (MPro) of SARS CoV-2 is another important enzyme translated by the virus, which is responsible for the maturation of itself as well as other crucial polyproteins, especially replicase polyproteins to form active replication complex [11–14]. Antiviral drugs against SARS-CoV-2 have been developed to block viral entry into host cells and inhibit subsequent viral RNA synthesis and replication or viral self-assembly [15]. Therefore, considering their enormous role in the viral replication cycle, conservancy and accessible active sites make them ideal targets for antiviral drug design [16].

Despite the substantial efforts to manage this pandemic, the lack of maintaining social distancing guidelines, the emergence of new variants almost daily with increased infectivity and transmissibility, the absence of effective therapeutics, and the potential downfall of vaccine efficacy [17] are crucial barriers to sustain the infection and mortality. Researchers and policymakers are prioritizing vaccination to reduce hospitalization and mortality rates. The continuous emergence of variants may facilitate viral reinfection and dodge the acquired immunity from vaccination. At this point of the pandemic, finding a potential therapeutic agent for COVID-19 demands urgency. Repurposing the approved antiviral drugs designed for RNA viruses whose safety and experiments or clinical trials document pharmacokinetics parameters seems a practical approach rather than the costly and time-consuming de-novo design [18, 19].

Therefore, to develop effective and safe treatment options to combat COVID-19, various clinical trials are undergoing to determine the potentiality of existing antivirals as anti-COVID-19 treatment options. Especially antivirals for SARS (Severe Acute Respiratory Syndrome), MERS (Middle East Respiratory Syndrome), Malaria, and HIV (Human Immunodeficiency Virus) are thoroughly inspected by conducting clinical trials across the globe [20]. Broad-spectrum antivirals such as Chloroquine, Hydroxychloroquine, and Iopinavir/Ritonavir are considered first-line drugs against COVID-19, while hydroxychloroquine plus azithromycin, oseltamivir, interferon, ribavirin, favipiravir, ivermectin, tocilizumab, sofosbuvir, and ozone therapy will be considered if first-line drugs failed [1, 20].

Remdesivir is the center of attention as a potential anti-COVID-19 drug after promising results in animal models and some trials. Gilead Sciences initially developed Remdesivir (also known as GS-5734), a mono-phosphoramidite prodrug of an adenosine analog [21], as a potential treatment for Ebola virus infection [22]. The FDA approved Remdesivir on October 22, 2020, to treat COVID-19 patients because it prevents viral replication in human nasal and bronchial airway epithelial cells [23] by interfering with RdRp and Mpro, two proteins required for viral replication [24–25]. Wang et al. showed that the condition of COVID-19 patients did not significantly improve when Remdesivir was administered [21]. Also, data from some studies about the adverse effects of Remdesivir after administration in hospitalized patients raised concerns about its clinical use [26]. Hence, further investigations are needed to point out the definite anti-COVID-19 activity of Remdesivir and determine the dose and other aspects of clinical administration. This study aims to repurpose Remdesivir as a potential and secure therapy option for COVID-19 by the computational drug design method to enhance the drug’s efficacy and safety. We have designed several new derivatives of Remidesivir marked by changing their functional groups and subsequently performed pharmacokinetic, molecular dynamics (MD) simulation, and molecular docking studies to estimate their drug-likeness to predict how effectively these derivatives can inhibit RdRp and MPro. However, further in-vitro/vivo tests might be required for stronger validation of the interaction mediated by the designed drug derivatives. Figure 1 depicts the entire strategy for developing remdesivir derivatives acting against SARS-CoV-2 RdRp and MPro.

**Figure 1.**
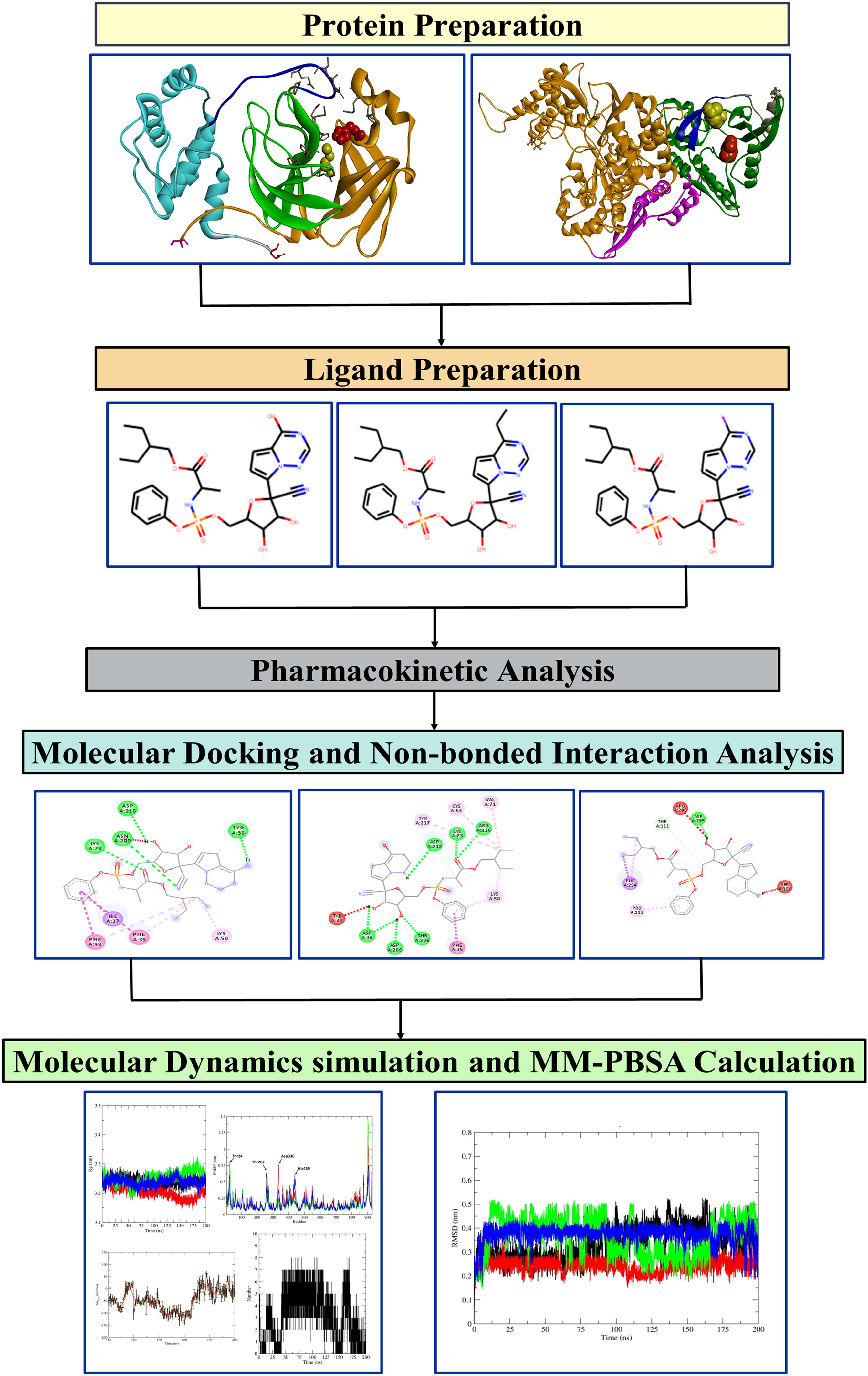
Schematic representation of methods for developing remdesivir derivatives against SARS-CoV-2 RdRp and MPro.

## 2. Methods

### 2.1 Ligands preparation

Remdesivir’s 3D structure was obtained from the structure data format (SDF) file of the PubChem online database (“Remdesivir | C27H35N6O8P - PubChem,” n.d.). Remdesivir’s structure was altered by adding functional groups with the chemical formulas C2H5, CF3, CH3, F, I, and OH in place of the NH2 group at position C-26. These functional groups are then designated as D-C2H5, D-CF3, D-CH3, D-F, I, and OH. The chem3D pro software minimized the ligands’ energy and their derivatives. The reduced structures of all ligands were recorded in SDF format for further investigation.

### 2.2 Target preparation for docking

A docking investigation of Remdesivir and its derivatives was performed against SARS-CoV-2 RdRp (PDB ID: 7BTF) and MPro (PDB ID: 6YB7) [27]. The structure of RdRp and MPro was downloaded in PDB format from the Protein Data Bank online database. For the molecular docking analysis in our work, a chain of targeted proteins was considered. Unwanted ions, ligands, functional groups, and water molecules were removed from the protein structure using the PYMOL program [28]. Swiss-PDB Viewer minimized energy in the improved protein structure, and the results were saved in PDB format [29].

### 2.3 Molecular Docking and Non-bond Interactions

Computer-assisted drug design relies heavily on molecular docking to forecast drug binding energy with target protein molecules [30]. Based on scoring, the best candidate from the library of chemicals is given upon successful docking and suggests a theory of how that ligand inhibits the target protein [31]. In this investigation, Remdesivir and its derivatives were molecularly docked using the Autodock Vina tool in PyRx software against the RdRp and then the MPro to identify prospective therapeutic compounds with the highest binding affinity [32]. The ligand-receptor complex with the lowest binding score exhibits the best interactions between the drug derivative and the target protein. The docked molecules were seen in the Biovia discovery studio visualizer, version 17.2, to show the drug-protein complex’s binding site and non-bond interactions [33]. For structure-based medication design in structural biology and pharmaceutical chemistry, it is helpful to recognize and quantify these non-bond interactions [34].

### 2.4 Pharmacokinetic parameters

Remdesivir’s pharmacokinetic characteristics and modifiers were assessed to determine their applicability and effectiveness as a treatment for RdRp and MPro. For Remdesivir and its derivatives, pharmacokinetic activity associated with drug absorption, distribution, metabolism, excretion, and toxicity (ADMET) was screened using MedChem Designer software and the new AdmetSAR online database [35]. SDF and the compounds’ simplified molecular-input line-entry system (SMILES) files were used to investigate the pharmacokinetic parameters.

### 2.5 Molecular dynamics simulation

#### 2.5.1 MD simulation and MM-PBSA calculations of RdRp to yield the best Remdesivir derivatives

The best docked remdesivir derivatives showing the most promising interactions with RdRp were selected for MD simulation analysis to validate whether the binding was stable. Through MD simulation and MM-PBSA calculations on the docked complex of D-CF3, D-I, D-OH, and remdesivir with RdRp and MPro, deeper insights into binding affinity and interactions were obtained. The 200 ns MD SIMULATION using Gromacs 2020.4 was conducted on the HPC cluster at the Bioinformatics Resources and Applications Facility (BRAF), C-DAC, Pune [36] MD simulation package. The missing residues of the loop segment were filled in by modeler 9.12 [37]. The ligand topologies were constructed from the CGenFF server using CHARMM General Force Field, whereas the protein topology was prepared using CHARMM-36 force field settings [38–40].

The ligand topologies were constructed from the CGenFF server using CHARMM General Force Field [39–40], whereas the protein topology was prepared using CHARMM-36 force field settings [38–39]. TIP3P water molecules [41] were first introduced as a solvent while holding a system in a dodecahedron unit cell. Next, the system was neutralized by adding Na+ counterions. Then, using the steepest descent minimization technique, the energy reduction phase was carried out to eliminate the steric conflicts until the threshold (Fmax 10 kJ/mol) was attained. A modified Berendsen thermostat [42] and a Berendsen barostat [43] were then used to equilibrate the interconnected systems under constant volume and temperature settings of 300 K for 100 ps each. All covalent bonds were regulated with the LINCS algorithm [45] throughout the 200 ns production phase MD simulation, which was carried out with the modified Berendsen thermostat and Parrinello-Rahman barostat [44]. The Particle Mesh Ewald technique (PME) was used to measure the long-range electrostatic interaction energies, and a cut-off of 12 Å was chosen [46]. Following the production phase MD simulation, the trajectories were examined for the radius of gyration (Rg), root mean square deviations (RMSD) in the backbone and ligand atoms, and root means square fluctuations (RMSF) in the side-chain atoms. Several hydrogen bonds were discovered. The binding free energy estimates were obtained using Poisson Boltzmann surface area continuum solvation (MM-PBSA) calculations on 500 MD snapshots isolated at 100 ps intervals between 150 ns and 200 ns [47–48].

#### 2.5.2 MD simulation and MM-GBSA calculations of MPro to yield the best Remdesivir derivatives

The best-docked derivatives with RdRp were further analyzed with MPro, and finally, the one showing the strongest binding affinity with MPro was selected for MD simulation analysis. The Desmond module of the Schrödinger LLC package was used in the MD simulation to investigate the alteration in protein structure within the solvent system [36]. Through Desmond’s System Builder panel, the ligand-protein complex was fixed using an orthorhombic periodic box soaked in solvent, with a minimum distance of 10 Å between the protein atoms and box edges. The solvent system was implemented using the single-point charge (SPC) water model [36–38]. The salt concentration was fixed to 0.15 M NaCl, which corresponded to the physiological system, and the charge of the constructed system was neutralized by adding Na+ and Cl-counterions. The solvated constructed system was decreased and relaxed using OPLS 2005 force field settings as the default protocol associated with Desmond [37]. MD simulations were performed using an isothermal, isobaric ensemble (NPT) with 300 K temperature, 1 atm pressure, and 200 ps thermostat relaxation time. The Coulombic interactions were calculated with a cut-off radius of 0.9 Å [39–41]. 2,000 trajectories were acquired during 200 ns of simulation. The Simulation Interaction Diagram (SID) tool was then used to investigate the MD simulation track. The generalized Born surface area (MM-GBSA) method and molecular mechanics were used to calculate the binding free energies of the ligand-protein complexes. Using the Python script (thermal mmgbsa.py), the average binding free energy (G Bind) based on MM-GBSA of the past 10 ns of simulation time using the VSGB solvation model linked to the OPLS3e force field was calculated [42].

## 3. Result and Discussion

### 3.1. Ligands preparation

Conformationally preferred functional groups, which are collections of linked atoms that can specify a parent molecule’s inherent reactivity and contribute to the overall properties of the molecule, are the cornerstone of contemporary medicinal chemistry. Halogenation is an effective approach to increasing the bioactivity of drugs so that it can play a pivotal role in drug development [49]. To further enhance the potential, iodine (-I) and fluorine (-F) was incorporated into the remdesivir parent drug. The D-I was modified with the iodine group replacing the -NH2 group at position 26C of remdesivir. In all vertebrates, iodine is required to generate thyroid hormone, so it is mostly used in thyroid hormone thyroxin drugs [50]. Although fluorine is a poor hydrogen bond acceptor and the most electronegative halogen element in the periodic table [51], it can take hydrogen bonds from H-bond donors [52]. In medicinal chemistry, fluorine presents interesting opportunities for enhancing the binding affinity of potential medication candidates. Trifluoromethyl (-CF3) [53] chemical groups are useful in current medication design because of these qualities [50]. The trifluoromethyl group also has strong electronegativity and hydrophobicity, which are useful in drug development to improve pharmacological activity [54]. Also, it can be linked with a wide range of organic compounds and is commonly used in the chemical and pharmaceutical sectors [55] [56]. Furthermore, lipophilicity is a significant compound feature that has attracted much attention in medicinal chemistry. It is connected to certain ADMET (Absorption, Distribution, Metabolism, Excretion, and Toxicity) factors and relates to the general “quality” of a molecule as a potential therapeutic candidate [53, 57]. In a systematic investigation of lipophilicity alterations caused by partial fluorination of n-alkyl groups connected at C3 of the indole unit, a distinctive lipophilicity pattern, CH3 >> CH2F = CHF2 CF3, appeared for terminally fluorinated n-propyl groups [58].

The trifluoromethyl group (-CF3) was integrated to position 26C of the remdesivir parent drug to replace the -H2 group in the modified drug derivative D-CF3. In modified drug derivatives D-CH3 and D-C2H5, the methyl (-CH3) and ethyl (-C2H5) groups were added to 26C of the remdesivir parent drug to replace the -NH2 group. The ortho effect, inductive effect, and conformational effect of alkyl groups (e.g., methyl and ethyl groups) can influence the physicochemical, pharmacodynamic, and pharmacokinetic properties of drugs. Furthermore, incorporating methyl into drug compounds can be used to create me-too drugs by finding new uses for old drugs [59]. Hydroxyl groups (-OH) form extended hydrogen bond networks in the target protein’s active site, enhancing affinity by several orders of magnitude. The polarized oxygen-hydrogen bond of hydroxyls facilitates hydrogen bond formation with suitable targets, such as functional groups or solvent molecules. The hydroxyl group was substituted for the –NH2 group at position 26C of remdesivir in the D-OH. Supplementary Figure S1 shows the two-dimensional structure of the parent drug remdesivir and its derivatives integrated with conformationally favored functional groups.

### 3.2. Target preparation

Coronaviruses produce a set of non-structural proteins from ORF1a and ORF1ab viral polyproteins. The key non-structural protein, NSP12, also called RdRp, is important in viral replication and transcription [60]. NSP12 constitutes an N-terminal β-hairpin (Asp29-Lys50), a nidovirus-specific N-terminal extension domain that forms a nidovirus RdRp-associated nucleotidyltransferase (NiRNA) (Asp60-Arg249), RdRp domain (Ser367-Phe920) and interfaces domain (Ala250-Arg365) (**Figure 2A**). Further, the RdRp domain makes the fingers, palm, and thumb regions [61]. RdRp was selected as a potential target against remdesivir and its modified derivatives since it is thought to be the major target for the approved drug remdesivir.

**Figure 2.**
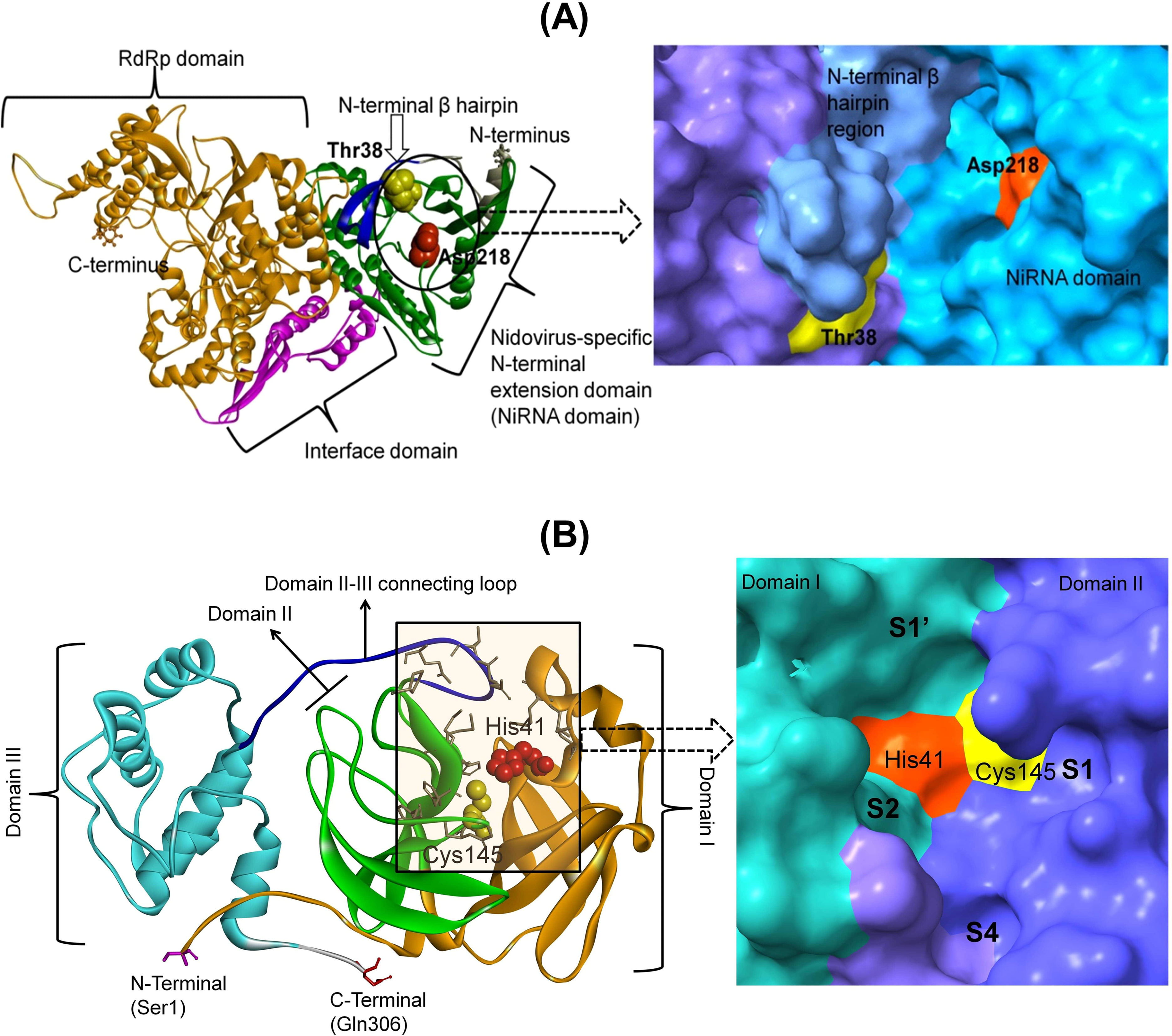
(A) The NSP12 structure shows the potential binding site. (The RdRp domain: orange, the interface domain: magenta, the NiRNA domain: green, and the β-hairpin: blue) (B) The SARS-COV-2 main protease structure shows a substrate binding site (left). (Domain I: orange, domain II: green, domain III: cyan, loop connecting domain II and domain III: blue. The substrate-binding site residues His41: red CPK, Cys145: yellow CPK and other residues are shown in stick representation). The surface representation with S1’, S1, S2, and S4 pockets and domains I and II are differently colored (right).

Furthermore, because MPro of SARS CoV-2 is now considered a promising therapeutic target for remdesivir, it was selected for further evaluation of the modified remdesivir derivatives in this study. The MPro is a homodimer with three domains (Domains I, II, and III). Domains I and II are made up of six antiparallel β-barrels, with residues 8-101 and 102-184, respectively, whereas domain III (residues 201-303) is formed by an antiparallel globular cluster of five helices that is linked to domain II by a lengthy loop region (residues 185-200). A Cys-His catalytic dyad in the gap between domains I and II, together with N-terminus residues 1 to 7, is critical in proteolytic activity [62–66]. The substrate-binding site in the cleft between domains I and II and the protomers between domains II and III are important in developing the substrate-binding site [64, 67–70]. Further, the substrate-binding cleft comprises 4 subsites, i.e., S1’, S1, S2, and S4 (**Figure 2B**) [71–72].

### 3.3. Analysis of binding affinity and non-bond interaction

Once the ligands (i.e., D-C2H5, D-CF3, D-CH3, D-F, D-I, D-OH) and the target proteins (i.e., RdRp and MPro) were prepared, they were subjected to molecular docking to retrieve the non-bond interactions between them. All the modified drug derivatives showed significantly higher binding affinity and non-bond interaction with RdRp than the remdesivir parent drug. The drug derivatives with the highest RdRp binding potential were examined further for their interaction with MPro. D-I, D-CF3, D-OH, D-CH3, and D-C2H5 all showed increased binding affinity of -9.5, -8.8, -8.9, -8.5, and -8.2 kcal/mol with RdRp, whereas remdesivir had a binding affinity of -8.0 kcal/mol. Furthermore, D-I, D-CF3, and D-OH had a higher binding affinity to MPro, with values of -6.6, -7.5, and -7.1 kcal/mol, respectively, than the remdesivir parent drug, which had a value of -7.0 kcal/mol. As a result, all modified drug derivatives appeared to bind with target proteins more strongly than the parent drug. These modified drug derivatives exhibited strong hydrogen and hydrophobic bond interactions with RdRp and MPro, respectively, as shown in Supplementary Figures S2 and S3. D-CF3 showed the strongest hydrogen and hydrophobic bond interactions with RdRp and MPro, as in Table 1. In an open conformational environment of protein structures, weak intermolecular interactions such as hydrogen bonding and hydrophobic interactions play crucial roles in stabilizing energetically-favored ligands. The binding affinity can be improved by adding conformationally favorable functional groups to the ligand-target interface’s active site. Stabilizing ligands at the target site are facilitated by hydrogen bonding and enhanced hydrophobic interactions, which also affect binding affinity and therapeutic efficacy [73].

**Figure 3.**
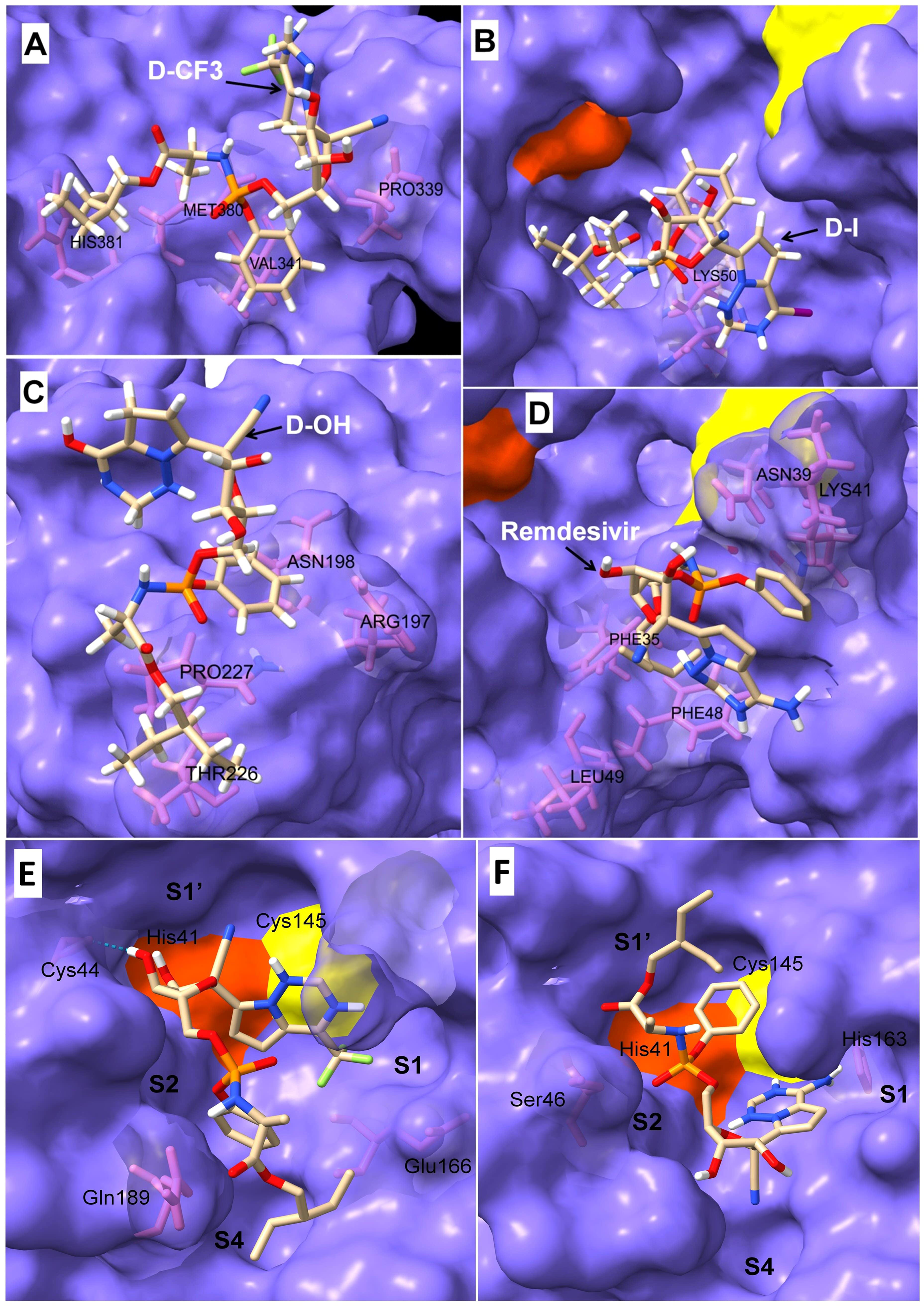
The protein-ligand interactions of RdRp with remdesivir derivatives (A) D-CF3, (B) D-I, (C) D-OH, and (D) remdesivir itself at the surface binding cleft of RdRp (Snapshots were taken at around 200 ns). The protein-ligand interactions at the substrate-binding cleft of MPro with (E) D-CF3 and (F) remdisivir parent drug.

**Table 1:**
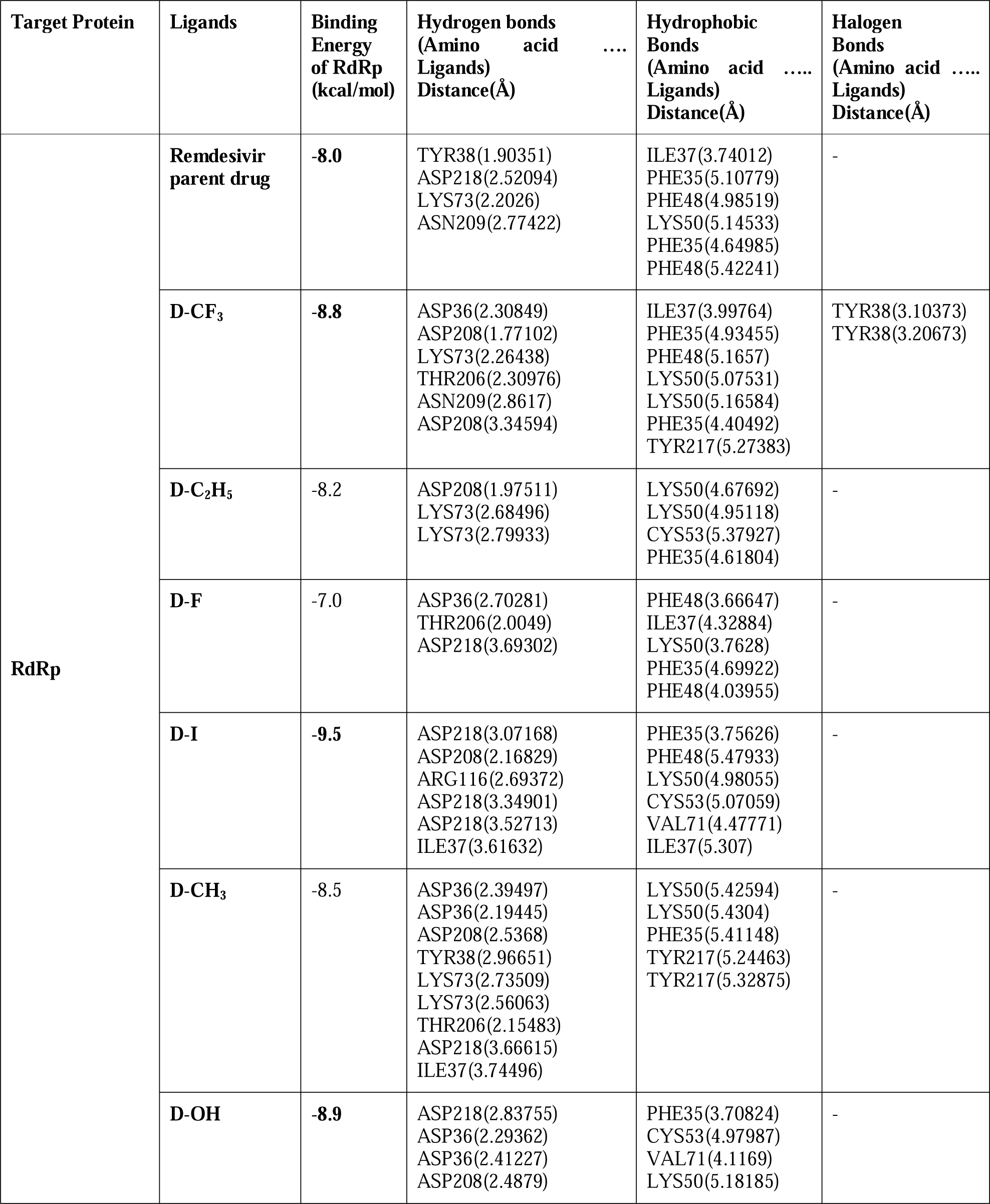
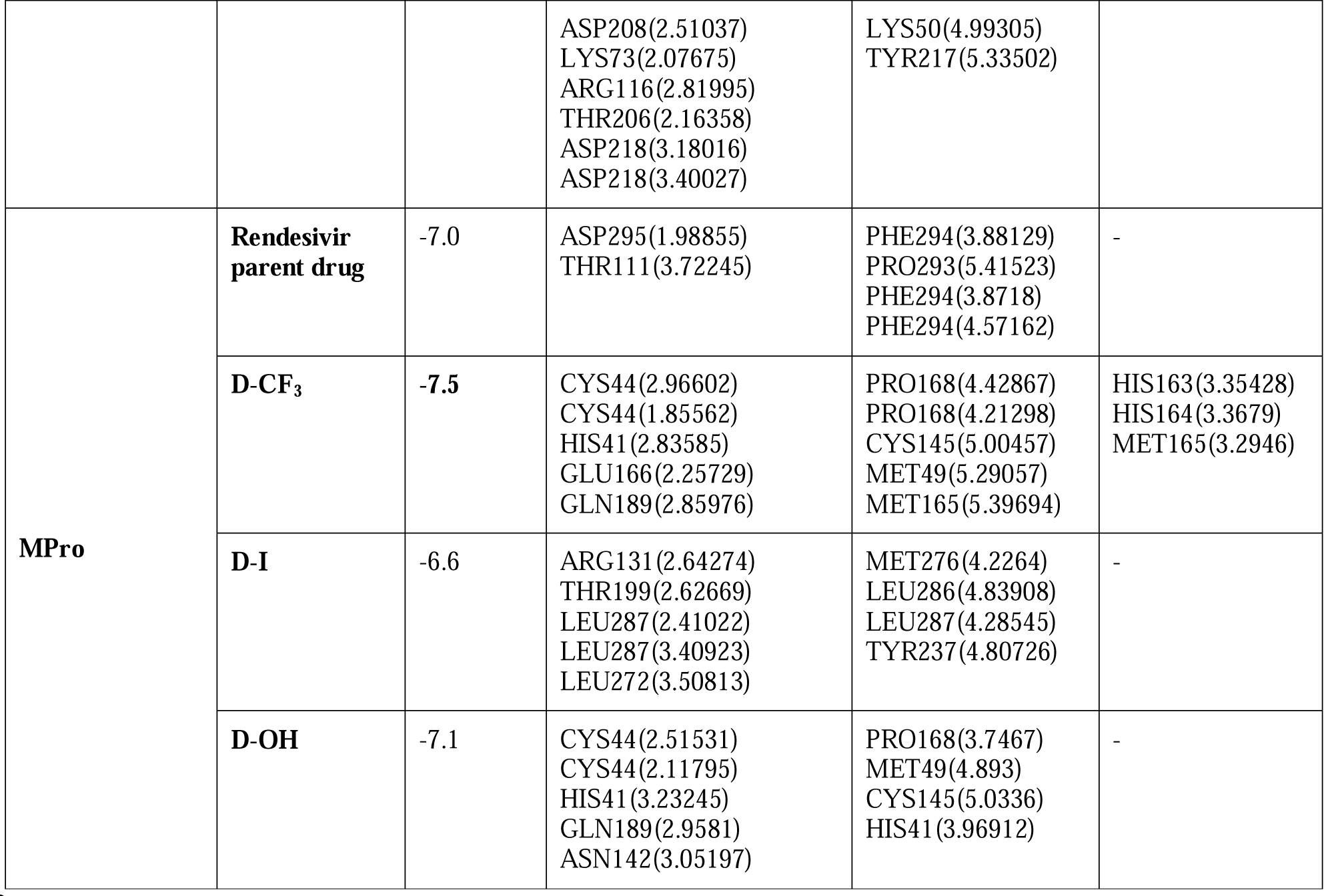
Binding energy and Non-bond interactions of Remdesivir and its derivatives against RdRp, generated via flexible docking.

### 3.4. Analysis of pharmacokinetic activity

The blood-brain barrier (BBB) prevents vital nutrients or medications from reaching the brain while shielding it from circulating toxins or bacteria that could cause illnesses. During the analysis of the pharmacokinetic activity of the remdesivir derivatives, they appeared to show favorable reactions with the BBB, indicating that they can pass through it. Low hydrogen-bonding potential, small size and molecular weight, and high lipophilicity are desired pharmacological qualities for bridging the BBB [74]. Human intestinal absorption (HIA) prediction has become extremely valuable as drug discovery processes have become more complicated. Intestinal permeability is measured using the Caco-2 permeability assay. The Caco-2 permeability assay from Cyprotex is based on a tried-and-true technique for calculating in vivo drug absorption by measuring the flow rate of a substance across polarised Caco-2 cell monolayers. Positive intestinal absorption scores and low caco-2 permeability are signs of adequate medication bioavailability [75–76]. Accordingly, all the remdesivir derivatives were predicted to have a high intestinal absorption potential. P-glycoprotein is an ATP (adenosine triphosphate)-binding cassette transporter that promotes multidrug resistance through the active efflux of different chemotherapeutic drugs. P-glycoprotein controls drug absorption and distribution in various organs, including the intestines and the brain. As a result, predicting P-glycoprotein-drug interactions is critical for evaluating drug pharmacokinetics and pharmacodynamics.

Positive P-glycoprotein inhibition scores can also guarantee the avoidance of the potential buildup of the remdesivir derivatives in the brain and their proper elimination [77]. The potassium ion (K+) channel encoded by the hERG (human ether-a-go-go-related gene) plays an important function in cardiac repolarization. Torsades de Pointes, a potentially fatal ventricular tachycardia, has been linked to drug-induced hERG inhibition. Drugs that inhibit the Human Ether-a-go-go Related Gene (hERG) can cause ventricular arrhythmia, which, in the worst-case scenario, can result in cardiac death [78–79]. Furthermore, the remdesivir derivatives were found to be non-carcinogenic and have negative hERG scores, indicating that the drugs could be safe for future use. Table 2 lists all parameters for determining pharmacokinetic activity retrieved from the updated version of AdmetSAR@LMMD.

**Table 2:**
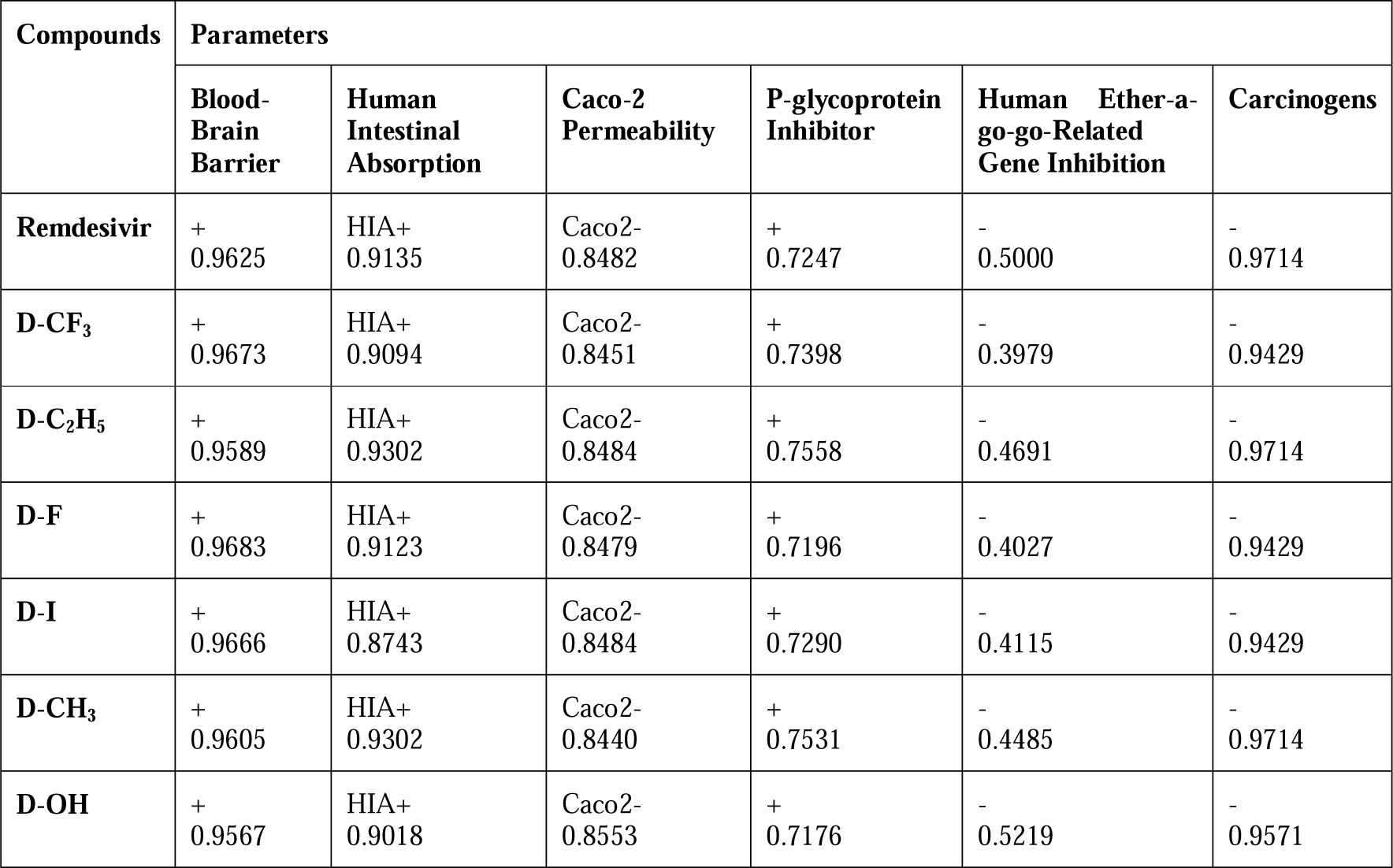
Remdesivir and its derivatives’ Selected pharmacokinetic parameters were obtained using AdmetSAR’s new version online database.

The pharmacokinetic activity of the remdesivir derivatives was further examined using MedChem Designer Software. The partition coefficient P (logP) between octanol and water (buffer), which represents the partition of the drug’s unionized (neutral) form, is the software’s definition of a drug’s lipophilicity. In contrast, logD describes the entire partition of the ionized and unionized forms [80]. All of the remdesivir derivatives examined in this study had logP values under 5, showing their hydrophilic character. Compounds with logP values [5 are lipophilic. A substance’s or compound’s capacity to permeate lipid-rich regions from aqueous solutions is known as lipophilicity [81]. Lipinski’s rule [82–83] states that a molecular weight of less than 500 Da, S+logp, and S+logD values [5, MlogP values [4.15, and a minimum of 5 hydrogen bond donors are all necessary for a molecular to pass through a biological membrane.

Furthermore, a logP (MLogP) value greater than 4.15 indicates that the molecule will be poorly absorbed [83]. All remdesivir derivatives had MlogP, S+logP, and S+logD values less than 5, indicating that they were hydrophilic and quickly absorbed and excreted. As a result, all evaluated properties indicated that the modified remdesivir derivatives are suitable for human use and could be administered without causing adverse side effects (**Table 3**).

**Table 3:** Pharmacokinetic properties of Remdesivir and its derivatives obtained from MedChem Designer Software.

| Compounds | Pharmacokinetic Parameters |  |  |  |  |
| --- | --- | --- | --- | --- | --- |
|  | MWt | MlogP | S+logP | S+logD | HBDH |
| **Remdesivir** | 602.588 | 0.634 | 1.597 | 1.597 | 5.000 |
| **D-CF<sub>3</sub>** | 655.572 | 1.424 | 2.905 | 2.905 | 3.000 |
| **D-C<sub>2</sub>H<sub>5</sub>** | 615.627 | 1.059 | 2.255 | 2.255 | 3.000 |
| **D-F** | 605.564 | 1.033 | 2.413 | 2.413 | 3.000 |
| **D-I** | 713.469 | 1.327 | 2.678 | 2.678 | 3.000 |
| **D-CH<sub>3</sub>** | 601.600 | 0.864 | 2.069 | 2.069 | 3.000 |
| **D-OH** | 603.573 | 0.634 | 1.611 | 1.581 | 4.000 |

### 3.5. Molecular dynamics simulation

In this study, a docked complex of D-CF3, D-I, D-OH, and remdesivir with RdRp was exposed to a 200 ns MD SIMULATION to capture stability and secondary structural changes in the RdRp structure and to acquire a deeper understanding of the binding mechanism of these ligands. The secondary structure of RdRp was found stable throughout the MD simulation, with some fluctuations in the loop region. Whether ligands remained bound in the binding pocket were analyzed by visual inspection of the trajectories at 0, 50, 100, 150, and 200 ns (**Supplementary Figures S4-S7**). All the ligands (i.e., remdesivir and its derivatives: D-I, D-OH, and D-CF_3_) remained bound at the binding pocket except on a few occasions during the MD SIMULATION (Figure 3A-D). The un-restrained production phase 200 ns MD SIMULATION showed that the ligands bind in the binding pocket differently than the one observed during the equilibrated condition.

The equilibrated system of docked complexes of MPro with remdesivir and D-CF3 was subjected to a 200 ns MD simulation to understand the binding mechanism better and determine the stability and potential secondary structural changes in the MPro structure. D-CF3 appeared to bind at the binding pocket during the MD simulation (Figure 3E-F). In this investigation, a duration of 200 ns was used, giving the MPro backbone atoms enough time to assume their complex configurations with the ligands. All the MD simulation trajectories were subjected to comparative analysis of RMSD, RMSF, contact map, and the percentage occupancies of the different types of interactions.

#### 3.5.1 Root mean square deviations (RMSD) analysis

The RMSD between corresponding atoms in two protein chains is a frequently used indicator of how similar two protein structures are. The RMSD reveals the overall stability of the protein-ligand combination in the atoms of the protein backbone and ligand, with lower values of RMSD indicating better stability (Reva, Finkelstein, Skolnick 1998). The RMSD in RdRp backbone atoms for systems with D-OH and D-CF3 bound has average values of 0.2881 and 0.3158 ns, respectively, which are lower than the RMSD for a system with bound remdesivir (**Supplementary Table S1**). In contrast, the average RMSD value for the system bound with D-I is slightly larger (Figure 4A-B). Prima-facie, these results of RMSD in RdRp backbone atoms suggest better stability of systems with D-CF3 and D-OH. The RMSD in atoms of D-I (0.2439 nm) is the lowest, while for other ligands, it is in the range of 0.3374 to 0.3751 nm. The iodo substituent is probably responsible for the D-I conformation’s stability. While the structures of D-CF3 and D-OH deviate probably to adopt stable conformations.

**Figure 4.**
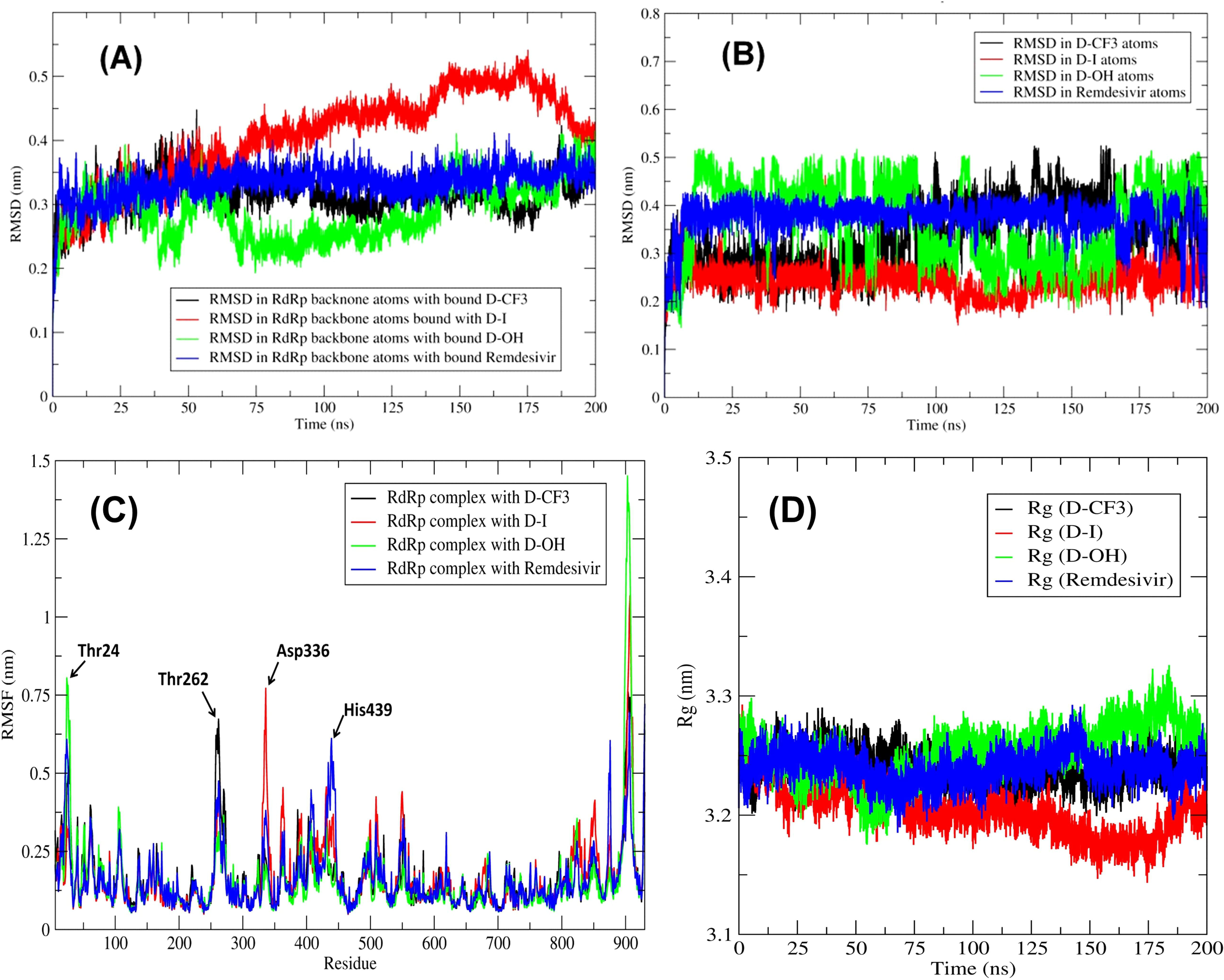
The root means square deviations (RMSD) evaluation. A) The RMSD in RdRp backbone atoms, B) The RMSD in ligand atoms, C) The root mean square fluctuation (RMSF) analysis, D) The radius of gyration (Rg) of RdRp bound with ligand

Interestingly, remdesivir conformation remains stable throughout the MD simulation with an average RMSD of 0.3751 nm (**Supplementary Table S1**). The residues interacting with the ligand might undergo major fluctuations. Such fluctuations in side-chain residue atoms were analyzed through the RMSF measurement.

Additionally, the stability, conformational behavior, and structural features of the protein-ligand complexes of MPro with remdesivir and D-CF3 were explained using MD simulations. For 200 ns simulations, the RMSD value for the C backbone was computed to evaluate the stability of remdesivir in a complex with MPro. The RMSD of the protein’s backbone ranged from 2.0 Å to 3.2 Å, with an average of 2.4 Å (**Figure 5A**). However, the RMSD of the remdesivir corresponding to the protein’s backbone fluctuated between 1.5 Å and 3.5 Å but was consistent with its structure, and it appeared to be quite stable, with lig-fit-prot deviations below 1.5 Å across the simulation time (**Figure 5B**).

**Figure 5.**
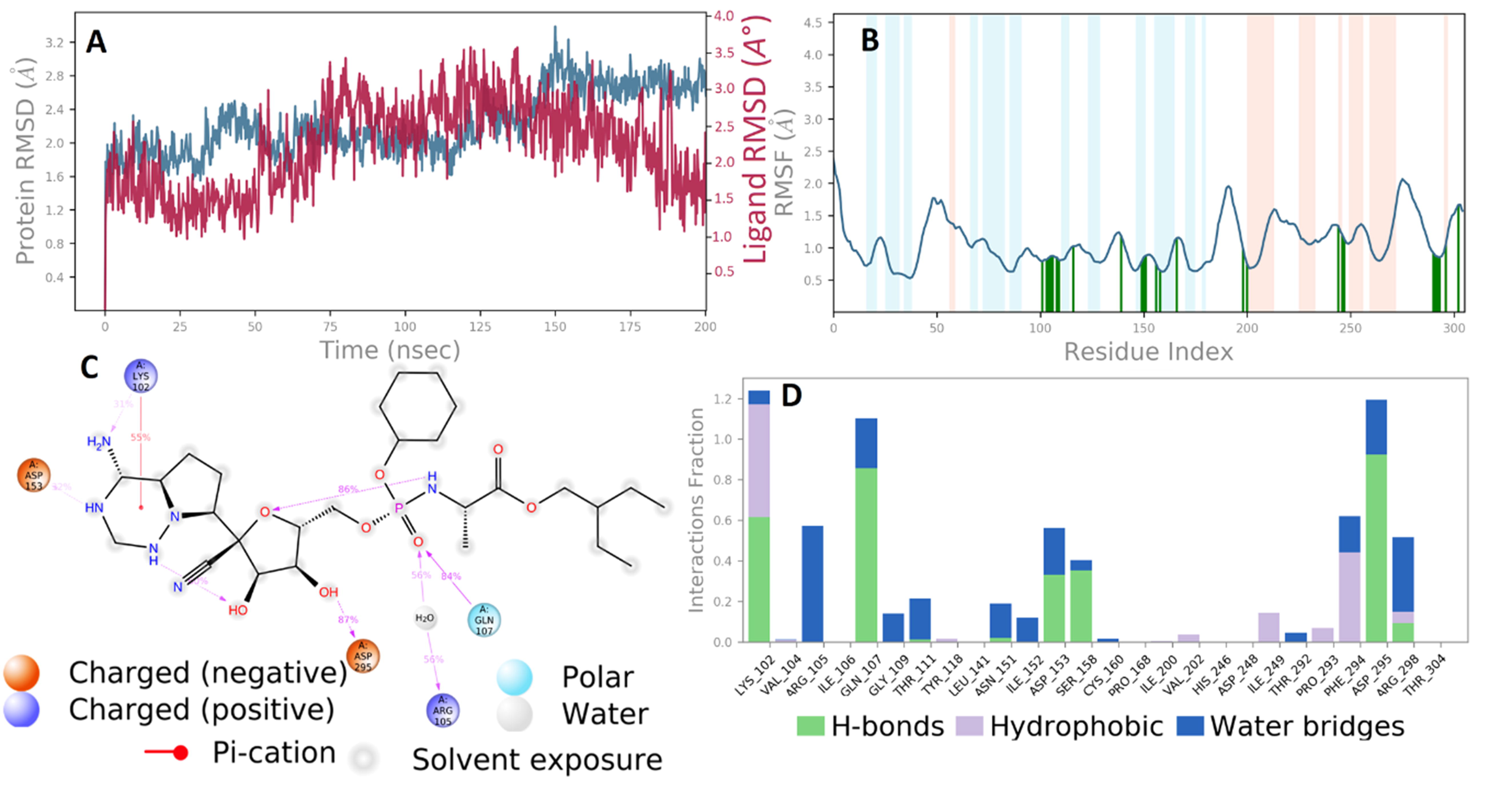
A study of the Remdesivir-MPro complex’s (A) RMSD using MD simulation (Protein RMSD is shown in grey while RMSD of remdesivir is shown in red) (B) Protein RMSF, (C) a 2D interaction diagram, and (D) a study of the MD trajectory’s protein-ligand contact.

On the other hand, simulation of the MPro-D-CF3 complex revealed that the MPro backbone had a maximum RMSD value of 3.6 Å, indicating that the protein complex remained stable for the simulation (**Figure 6A**). The RMSD of the D-CF3 in complex with MPro remained between 2.4 Å and 5.6 Å till 175ns. Afterward, insignificant RMSD fluctuation was noticed and remained stable for the rest of the simulation (**Figure 6B**). It is important to note that the MPro protein’s RMSD fluctuations were close to or below the permitted limit of 3 Å throughout the simulation [84]. As a result, the observations show that the binding of D-CF3 and remdesivir did not appreciably change the overall structures of the RdRp and MPro, and the protein-ligand complexes remained largely stable during the simulation.

**Figure 6.**
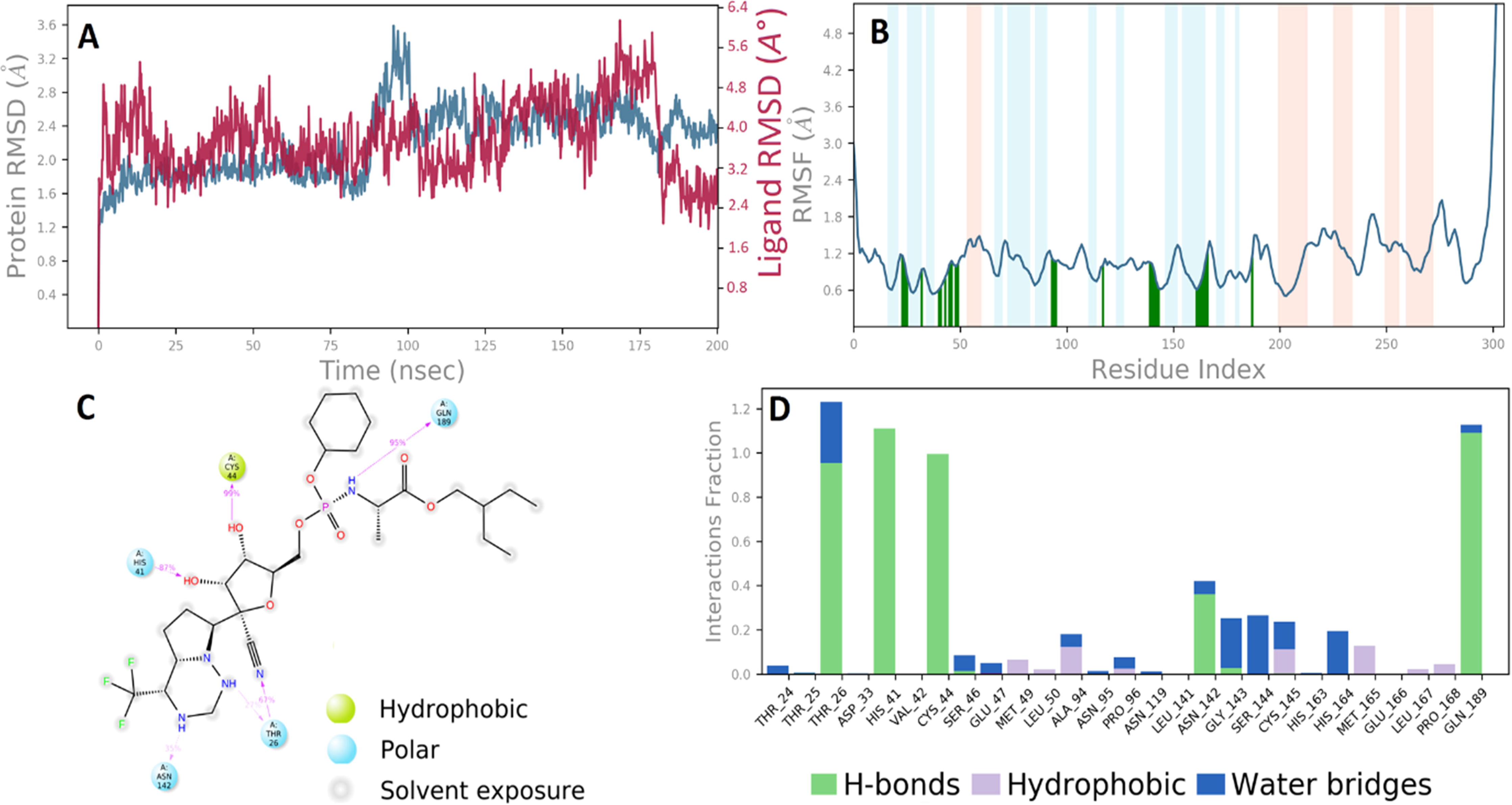
The analysis of the D-CF3-MPro complex’s (A) RMSD using MD simulation (Protein RMSD is shown in grey while RMSD of D-CF3 is shown in red) (B) Protein RMSF, (C) a 2D interaction diagram, and (D) a study of the MD trajectory’s protein-ligand contact.

#### 3.5.2 Root mean square fluctuations (RMSF) analysis

Protein-ligand complexes’ flexible areas can be discovered by analyzing RMSF plots of the complexes. White represents the loop region, while red and blue represent additional structural elements such as the -helical and -strand regions. They change less than loop regions because -helical and -strand parts are usually stiffer than the unstructured component of the protein. The target protein’s main chain and active site atoms vary slightly. There has not been much conformational shift in that situation, indicating that the ligand is tightly bound inside the target protein binding pocket [85–87].

In the RMSF study of RdRp, a few residues of RdRp interacting with the ligands seemed to be fluctuating to a larger extent. Figure 4C shows the results of the RMSF measurement. The residues from 250 to 460 show major fluctuations. This part belongs to the interface domain and the RdRp domain. The non-terminal residue Thr262 seems to undergo the largest fluctuation in the system with D-CF3, while Asp336, Thr24, and His 439 similarly showed the largest fluctuation in a system with D-I, D-OH, and remdesivir, respectively. The equilibrated system of RdRp with D-CF3 showed hydrogen bonds with Thr206, Tyr38, Asp208, Asn209, and Lys73. None of these hydrogen bonds remained stable, and new hydrogen bonds with Leu49 standing out were seen during the MD SIMULATION **(Supplementary Figure S4).** While in the case of initial equilibrated system of RdRp with D-I showed hydrogen bonds with Tyr38, Asp36, Thr206, Asp208, Asn209, and Arg116. These hydrogen connections, however, were broken, and at about 50 ns, a new hydrogen bond was created with the residues Thr51 and Thr76. In the final 25 nanoseconds of the MD simulation, residues Thr76 and Lys50 displayed hydrogen bonding **(Supplementary Figure S5)**. The equilibrated system of RdRp with D-OH showed hydrogen bonds with Asp36, Thr206, Tyr38, Asp208, Asp218, Gly220, and Lys73. However, new hydrogen bonds between Asn198 and Glu84 were created after about 100 nanoseconds of MD simulation **(Supplementary Figure S6).** The equilibrated system with remdesivir showed hydrogen bonds with Asn209, Asp208, Thr206, and Tyr38. However, all of these hydrogen bonds disintegrate at around 50 ns, and new hydrogen bonds with Gly712, Asp711, and Leu49 form; these new hydrogen bonds last for 100 ns and are hence stable. Then, in the final stages of the MD simulation, additional hydrogen bonds were created with the residues Thr51 and Asn39 **(Supplementary Figure S7).** The RMSF plot shows that the majority of these residues were oscillating.

Remdesivir made contact with 26 amino acids of the remdesivir-MPro complex in the RMSF plot, including Lys102, Val104, Arg105, Ile106, Gln107, Gly109, Thr111, Tyr118, Leu141, Asn151, Ile152, Asp153, Ser158, Cys160, Pro168, Ile200, Val202, His246, Asp248, Ile249, Thr292, Pro293, Phe294, Asp295, Arg298, and Thr304. All interacting residues have an RMSF value of less than 2.0 Å, denoted by green vertical bars (**Figure 5B**). Further, in the D-CF3-MPro complex, D-CF3 interacted with 27 MPro amino acids, including Thr24, Thr25, Thr26, Asp33, His41, Val42, Cys44, Ser46, Glu47, Met49, Leu50, Ala94, Asn95, Pro96, Asn119, Leu141, Asn142, Gly143, Ser144, Cys145, His163, His164 Met165, Glu166, Leu167, Pro168 (**Figure 6B**). The RMSF values of most residues are [2 Å, except for the loop regions and N terminus, indicating that during the simulation, the residue structure is quite stable (Figure 5B and 6B). The RMSF figures above demonstrate that the MPro residues bound to D-CF3 remained constant throughout the run.

#### 3.5.3 The radius of gyration (Rg) analysis

The radius of gyration (Rg) analysis measures the system’s compactness (Shawon et al., 2018). The results of the total Rg of RdRp bound with ligands are shown in Figure 4D. The results of the comprehensive Rg analysis suggest that the RdRp structure is compact with few deviations, presumably originating from the flexibility in the loop regions. The system with D-I has the lowest total Rg of 3.2047 nm, while the systems with D-CF3, D-OH, and remdesivir have a total Rg of 3.2407, 3.2534, and 3.2396 nm, respectively. The slightly higher values for total Rg for these systems suggest the conformational changes in protein structure probably in loop regions to adopt the respective ligands in the surface binding cleft. The stability of the lead compounds in the SARS CoV-2 MPro binding pockets throughout a 200 ns simulation was also demonstrated by looking at the Rg property. To determine how stretched a ligand is, use the Rg parameter, which corresponds to its primary moment of inertia (Figure 7). The average Rg values for the lead compounds D-CF3 and Remdesivir in complexes with MPro were 4.87 [0.19 Å and 4.71 [0.12 Å, respectively. There were no discernible alterations, according to the Rg study. These constant values exhibited a consistent pattern.

**Figure 7.**
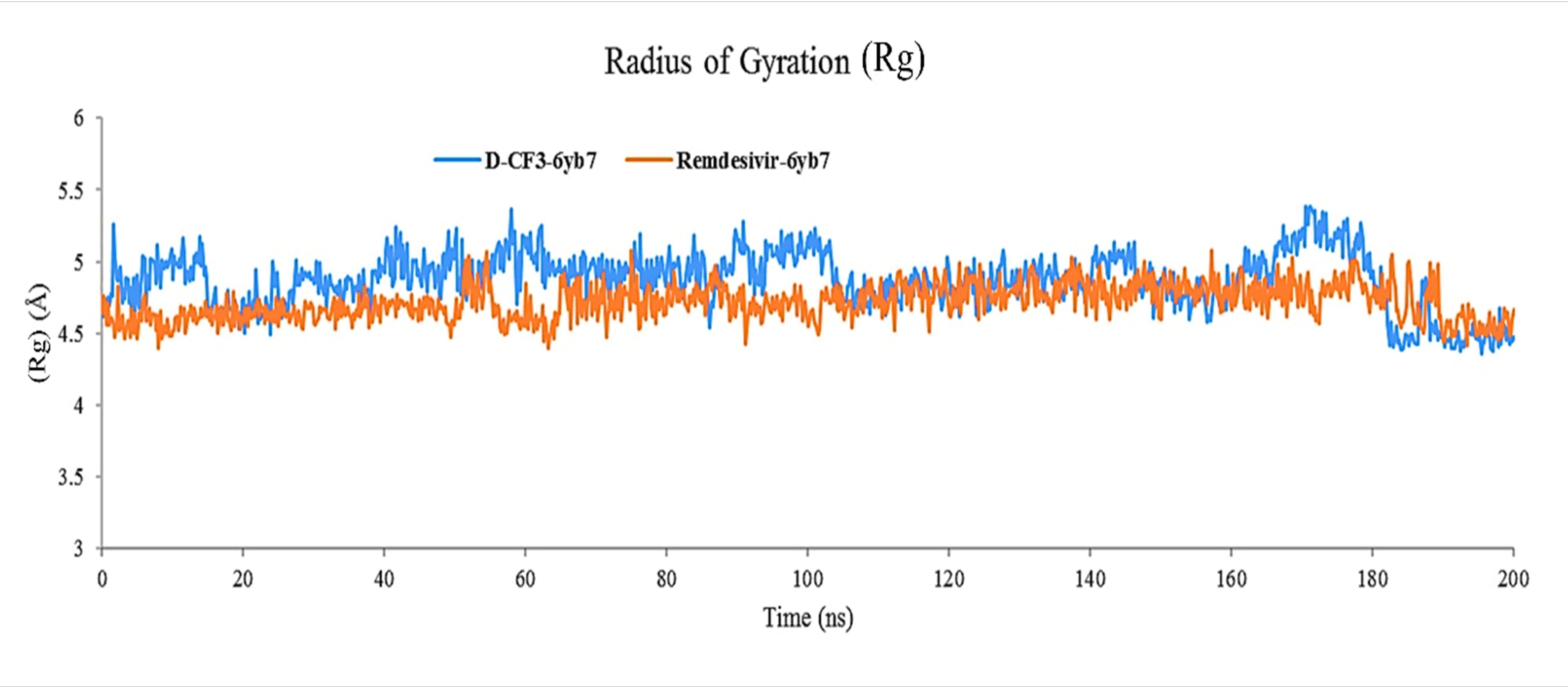
The radius of gyration (Rg) graph of a simulated complex of MPro with remdesivir and D-CF3 at 200 ns simulation time.

#### 3.5.4 Hydrogen bond interaction analysis

Non-bonded interactions, including hydrogen bonds, are crucial for stabilizing the system and determining the ligands’ propensity for binding [88]. On average, 2 hydrogen bonds were found forming between RdRp and D-CF3 (**Supplementary Figure S8**). However, no hydrogen bond is formed between 75-110 ns and around 140-160 ns. The visual inspection of snapshots revealed that the ligand D-CF3 moved out of the binding pocket during these time intervals. The ligand D-I showed around 3 consistently formed hydrogen bonds throughout the MD SIMULATION. The system with D-OH showed maximum 4 hydrogen bonds being formed. However, no hydrogen bonds were created throughout the 50-100 ns MD simulation period, and the ligand was observed leaving the binding pocket. Maximum hydrogen bonds that might form in the system with remdesivir were 6; however, on average, only 3 hydrogen bonds were regularly seen to form.

Furthermore, the hydrogen bond % occupancy results (**Supplementary Table S2**) suggest that D-CF3 forms a consistent hydrogen bond with a % occupancy of 11.8 % with Leu49. The ligand D-I forms a hydrogen bond with the highest % occupancy of 32.5 % with Arg74, while Asp221 and Thr76 form hydrogen bonds with a % occupancy of more than 20 % occupancy. In the case of ligand D-OH, the hydrogen bond with a % occupancy of 8.3 % was formed with residue Asn198. The remdesivir forms the hydrogen bond with a % occupancy of 63% with Leu49, while the residues Gly712 and Lys41 form the hydrogen bond with a % occupancy of more than 41%. The results suggest that ligands D-I and remdesivir could form stable hydrogen bonds.

On the other hand, protein-ligand contact mapping was conducted to explore the hydrogen bond interaction of remdesivir and the selected derivative: D-CF3, with MPro. The analysis showed that Remdesivir binding to the MPro involves hydrophobic interaction with Lys102, Val202, Ile249, Pro293, Phe294, and Arg298; polar interaction with Gln107, positively charged interaction with Lys102 and negatively charged interaction with Asp153 and Asp295 (**Figure 5D**). Additionally, the lead chemical D-CF3 showed more than 60% hydrogen bond interactions with the residues Thr26, His41, Cys44, and Gln189. Catalytic-colored residue His41 accounts for 87% of the hydrogen bond interaction with D-CF3 out of the group. During the MD simulation, the Met49, Leu50, Ala94, Cys145, and Met165 displayed hydrophobic interactions (**Figure 6D**). Most of the contacts between the MPro and D-CF3 seen during docking remained after the MD simulation, suggesting that the predicted binding mode is stable.

#### 3.5.5 MM-PBSA and MM-GBSA calculations

Binding affinity evaluation is one of the more accurate evaluations in determining the ligand’s ability to occupy the binding cavity under a simulated environment favorably. In the MM-PBSA calculations, the Adaptive Poisson-Boltzmann Solver (APBS) method was used to solve the continuum electrostatic equations of the system under study, and various energy terms, including van der Waal energy, electrostatic energy, polar solvation energy, SASA energy, and binding energy ([G_bind_), were estimated. [89]. The RdRp structure has more than 900 residues, which could increase the computational cost of the MM-PBSA calculations. For this, the last 500 photos that were isolated at intervals of 100 ps between 150 and 200 ns were used in the MM-PBSA calculations. Both Supplementary Table S3 and Supplementary Figure S9 provide the findings of the MM-PBSA computations. The most popular ligand is D-CF3, which has a binding free energy of -57.766 kJ.mol^-1^. Although the hydrogen bond analysis and visual inspection of all the snapshots suggested that it moved out of the binding pocket on a few occasions, it has the least polar solvation energy and reasonably favorable van der Waals and electrostatic energies compared to other ligands. In the case of ligand D-I, though it showed a better number of hydrogen bond interactions, the binding free energy is the least. The polar solvation energy for this ligand is the highest among all the ligands, which could be the reason for its higher binding energy.

Similarly, the ligand D-OH has slightly larger polar solvation energy and larger van der Waals and electrostatic energies than D-CF3. It has almost similar binding free energy as that of ligand D-I. Remdesivir has a binding free energy of -45.952 kJ.mol^-1^, which is higher than the ligand D-CF3, possibly due to higher polar solvation energy. Altogether, the ligand D-CF3 has the best binding free energy amongst all the ligands and could bind to RdRp with better affinity.

The post-dynamic MM-GBSA analysis of binding free energy ([G_Bind_) calculation for MPro complexes with remdesivir and D-CF3 was performed with the creation of 900-1001 frames having a 10-step sampling size to assess the binding association between the MPro and D-CF3 and remdesivir. A total of 11 frames were processed and analyzed throughout the 200 ns MD simulation data of lead compound in complex with the SARS CoV-2 main protease revealed by the dynamics studies. Supplementary Table S4 shows the contributions of all parameters to the binding free energy, demonstrating that the overall contributions of Coulombic, H-bond, Lipo, and vdW interactions significantly impact ΔG _Bind_. The average binding free energy ΔG _Bind_ of the complex D-CF3 and remdesivir in complex with the MPro was found to be -72.48 ± 3.46 kcal/mol, and -46.50 ± 3.96 kcal/mol, respectively. A lower number suggests a higher binding affinity because the MM/GBSA binding energies are estimations of binding free energies (Rastelli et al., 2010). Compared to the remdesivir complex MM-GBSA calculations, the D-CF3 complex revealed better binding free energy scores.

## 4. Conclusion

The COVID-19 pandemic is becoming outrageous day by day. While witnessing the resurrection of infections and death tolls, people hope to see an end to this pandemic. This scenario seems to continue shortly because of the frequent mutations in the SARS-CoV-2 genome, enabling the virus to be deadlier. The recent emerging Omicron variant of SARS-CoV-2 has provoked the almost 2-year-old COVID-19 pandemic’s seemingly everlasting burning. Healthcare providers and researchers have seen relentless efforts to limit infection by developing therapeutics and administering vaccines to people. Also, several clinical trials of FDA-approved drugs are in place to assess their applicability to treating COVID-19, but the initial findings of these trials are unsatisfactory. In our present study, we computationally designed derivatives of only FDA-approved drugs for COVID-19 to propose promising drug candidates without adverse side effects by replacing functional groups. We targeted two NSP proteins, namely RdRP and MPro, and assessed the inhibitory potential of our designed derivatives by molecular docking and dynamics simulation as well as pharmacokinetic parameters to find their drug-likeness. Data from our study revealed that our designed derivatives can strongly inhibit RdRP and MPro than the parent remdesivir and can be administered for treating COVID-19-infected patients without any potential side effects.

## Declarations

### Ethics approval and consent to participate

Not Applicable

### Consent for publication

Not Applicable

### Availability of data and material

All the data generated during the experiment are provided in the manuscript/supplementary material.

### Authors’ contributions

MS, YA, and KAA conceived the study. AM, KAA, CZ, and MS designed the study. AM, SZ, MFS, RBP, IA, TAH, RP, DM, MNS, NN, HP, JZ, MS, and KAA wrote the draft manuscript. TAH, MIH, MAH, ZAJ, KNU, SRS, MLK, MX, YA, CZ, ASC, and AM did the revisions. CZ supervised the study. All authors approved the manuscript submission.

## Acknowledgment

We take this opportunity to express our sincere gratitude to all individuals who have offered us support throughout our research endeavor. We extend our heartfelt appreciation to the esteemed faculty members of the Department of Genetic Engineering and Biotechnology at the University of Chittagong and Department of Biochemistry and Molecular Biology at Bangladesh Agricultural University for their valuable contributions to our education and research. Furthermore, we wish to acknowledge the support and encouragement extended to us by Dr. Naznin Naher Islam and Dr. Adnan Mannan during our research journey. We would like to dedicate this study to all aspiring student researchers in Bangladesh who are passionate about science and research, and whose contributions to the field will undoubtedly be significant. We are grateful to all our supporters for their contributions and encouragement during our research endeavor.

